## Supplementary material for "Sonogenetic control of mammalian cells using exogenous Transient Receptor Potential A1 channels": This file includes Extended Data Figures S1-13, Extended Data Tables S1-2 and Legends for Videos S1-S6.

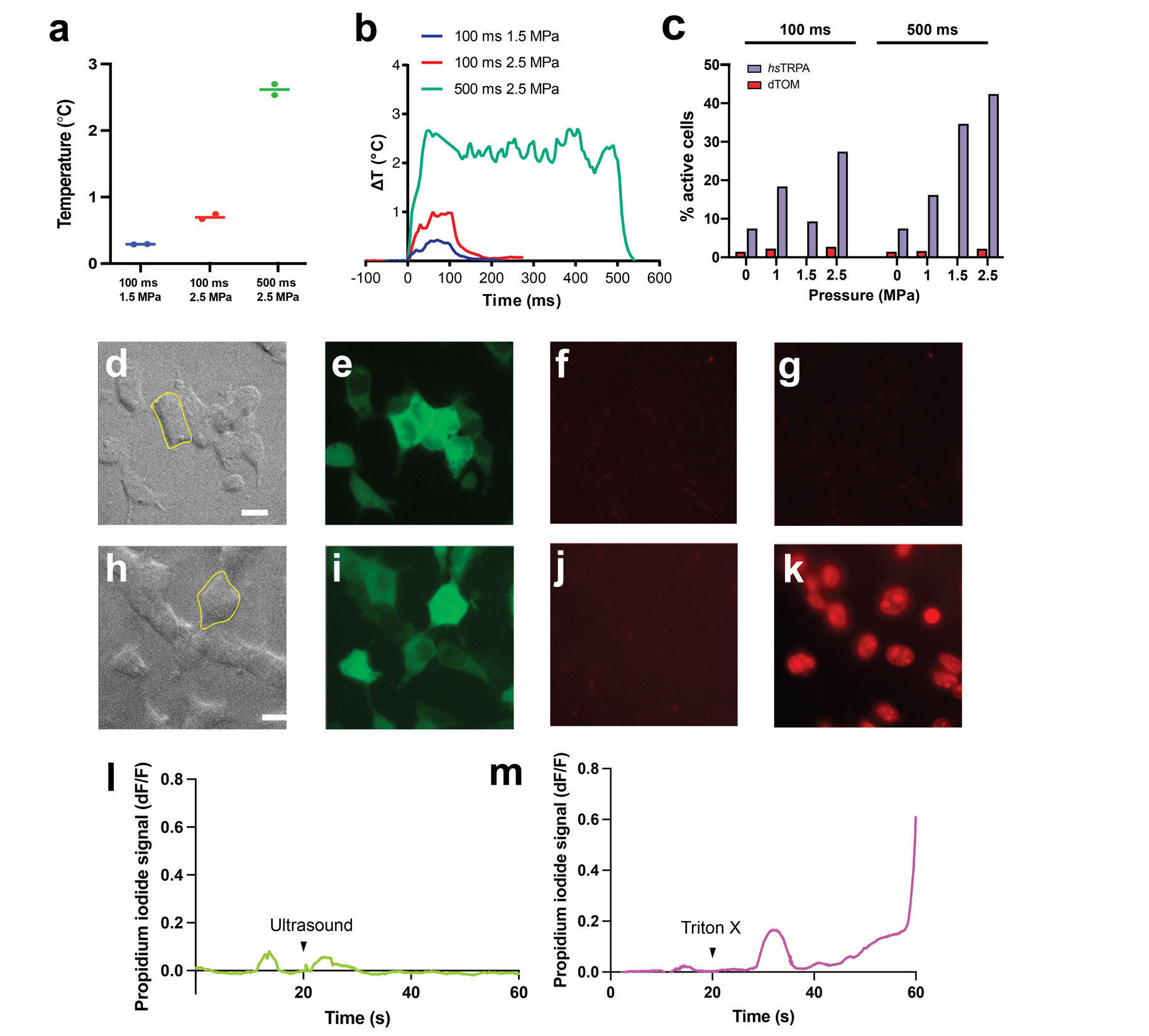

**Extended Data Fig. S1. Safety profile of 6.91MHz ultrasound stimulation in HEK cells. a**, Plot showing maximum temperature increases under different ultrasound stimulation parameters. n = 3 assays/condition. **b**, Time series of ultrasound-evoked temperature changes in the cell culture dish during stimulation. **c**, % active hsTRPA1 and dTom transfected cells in response to ultrasound at different pressure and durations. n = 3 coverslips/condition**. d,** Image showing bright field (BF) image for GCaMP6f-HEK cells, and the corresponding GFP channel (**e**) and propidium iodide channel (**f**) before ultrasound stimulation. Multiple trials with ultrasound stimulation at 2.5MPa 100ms had no effect on the intracellular levels of propidium iodide (**g**) n = 3 stims. **h,** Image showing HEK cells used for the positive control, including GFP channel (**i**) and propidium iodide channel before treatment (**j**). Addition of 0.1% Triton-X induced a significant increase of intracellular propidium iodide (**k**). Time course for the propidium iodide signal for an ultrasound stimulated cell highlighted in **d,** shown in **l,** and for a cell treated with propidium iodide highlighted in **h,** shown in **m.** Scale bar, 20 µm.

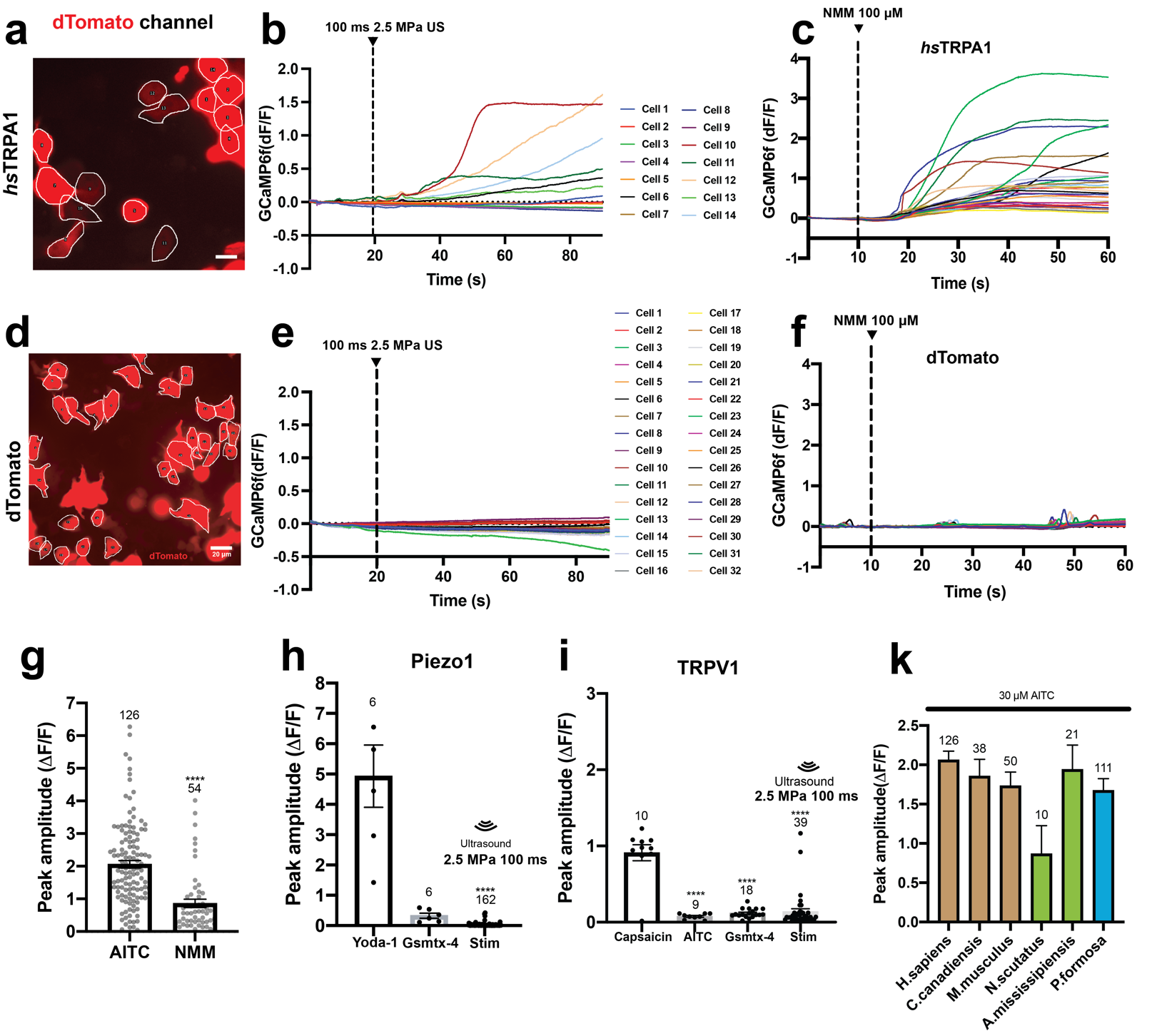

**Extended Data Fig. S2. Characterization of TRPA1 calcium responses in HEK cells. a**, Image showing dTom+ ROIs in HEK cells expressing hsTRPA1 and change in GCaMP fluorescence upon **b,** ultrasound stimulation or **c,** application of NMM in individual cells. **d**, Image showing HEK cells expressing dTom control and change in GCaMP fluorescence upon **e,** ultrasound stimulation in individual cells or **f,** application of NMM in individual cells. **g,** HEK cells expressing hsTRPA1 respond to TRPA1 agonists, N-methyl maleimide (NMM, 100 μM). and allyl isothiocyanate (AITC 33 μM). n = 3 coverslips/condition. **h**, HEK cells expressing mouse-Piezo1 respond to yoda-1(10 μM), but not GsMTx-4-4 or ultrasound. n = 3 coverslips/condition. **i**, HEK cells expressing human-TRPV1 respond to capsaicin (3 μM), but not AITC, GsMTx-4-4 or ultrasound. n = 3 coverslips/condition. Number of cells analyzed is shown on each bar. ****p<0.0001, by Kruskal-Wallis rank test and Dunn’s test for multiple comparisons. Scale bar, 20 µm. **k**, Response to AITC in HEK cells expressing TRPA1 from tested species.

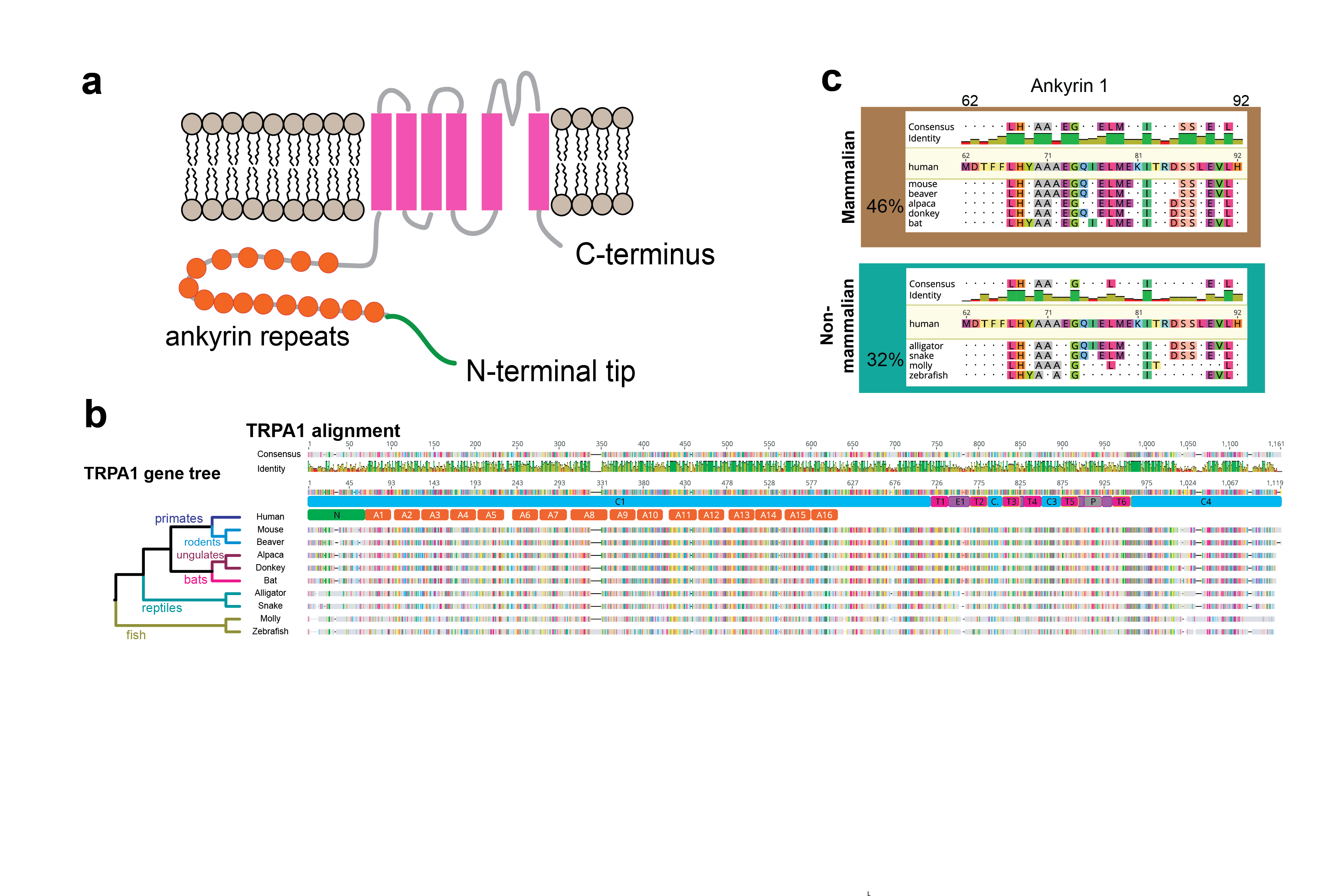

**Extended Data Fig. S3. TRPA1 sequence alignment across homologs tested for ultrasound sensitivity. a,** Schematic of TRPA1 showing the N-terminal region (green), 16 ankyrin repeats (orange) and the 6 transmembrane domains (pink). Mammalian and non-mammalian alignments of TRPA1 homologs tested for ultrasound sensitivity, depicting different domains and %identity compared to hsTRPA1 for the whole protein **b**, and for Ankyrin 1 **c**. % indicates % identity between 65% consensus sequence and *hs*TRPA1.

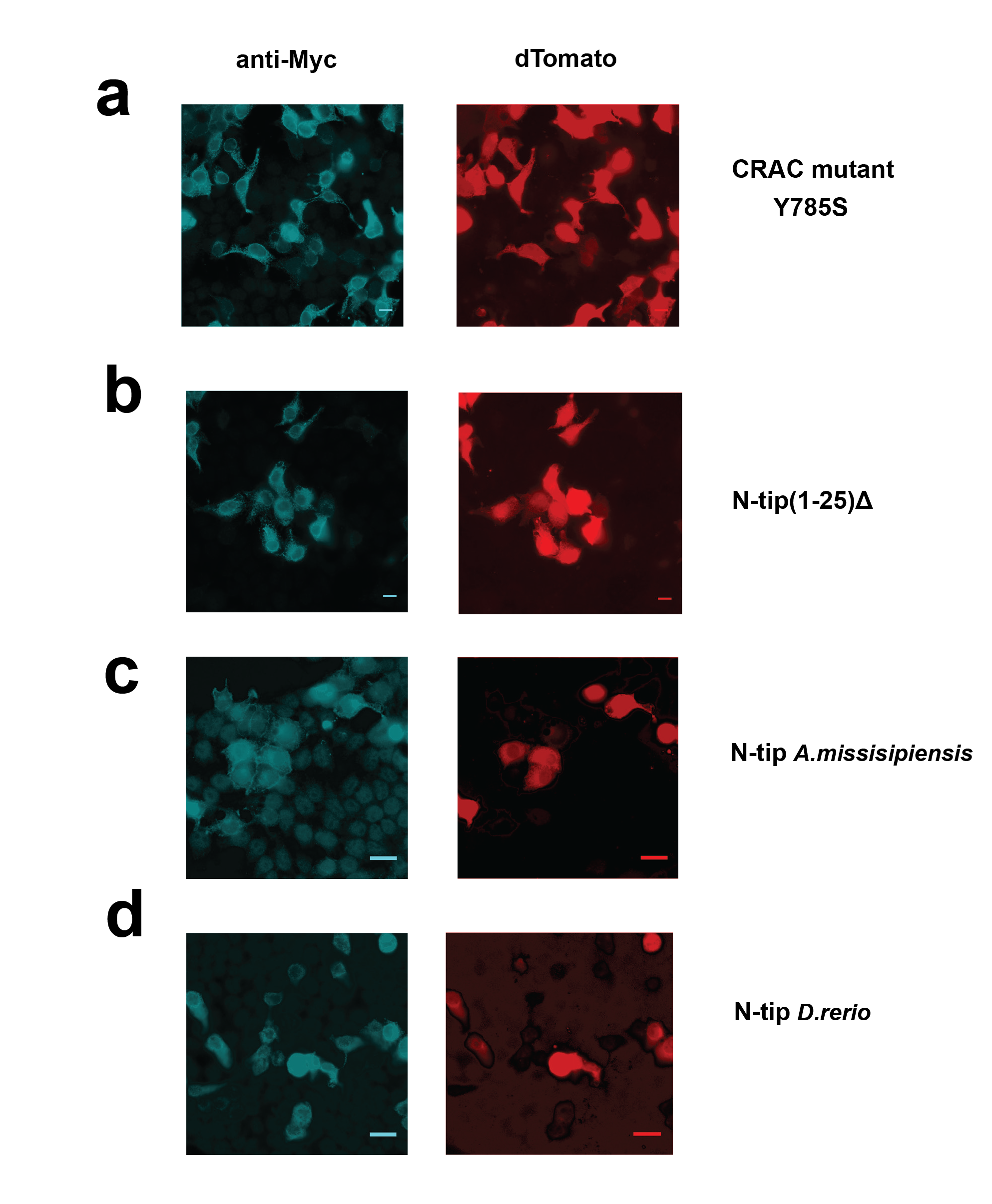

**Extended Data Fig. S4. Expression of TRPA1 mutants in HEK293T cells.** Immunohistochemistry showing expression and correct trafficking of myc-tagged TRPA1 constructs with **a,** CRAC Y785S mutation, **b,** N-terminal tip (1-25) deletion, **c,** *am*TRPA1 N-terminal tip swapped into *hs*TRPA1 and **d,** *dr*TRPA1 N-terminal tip swapped into *hs*TRPA1.

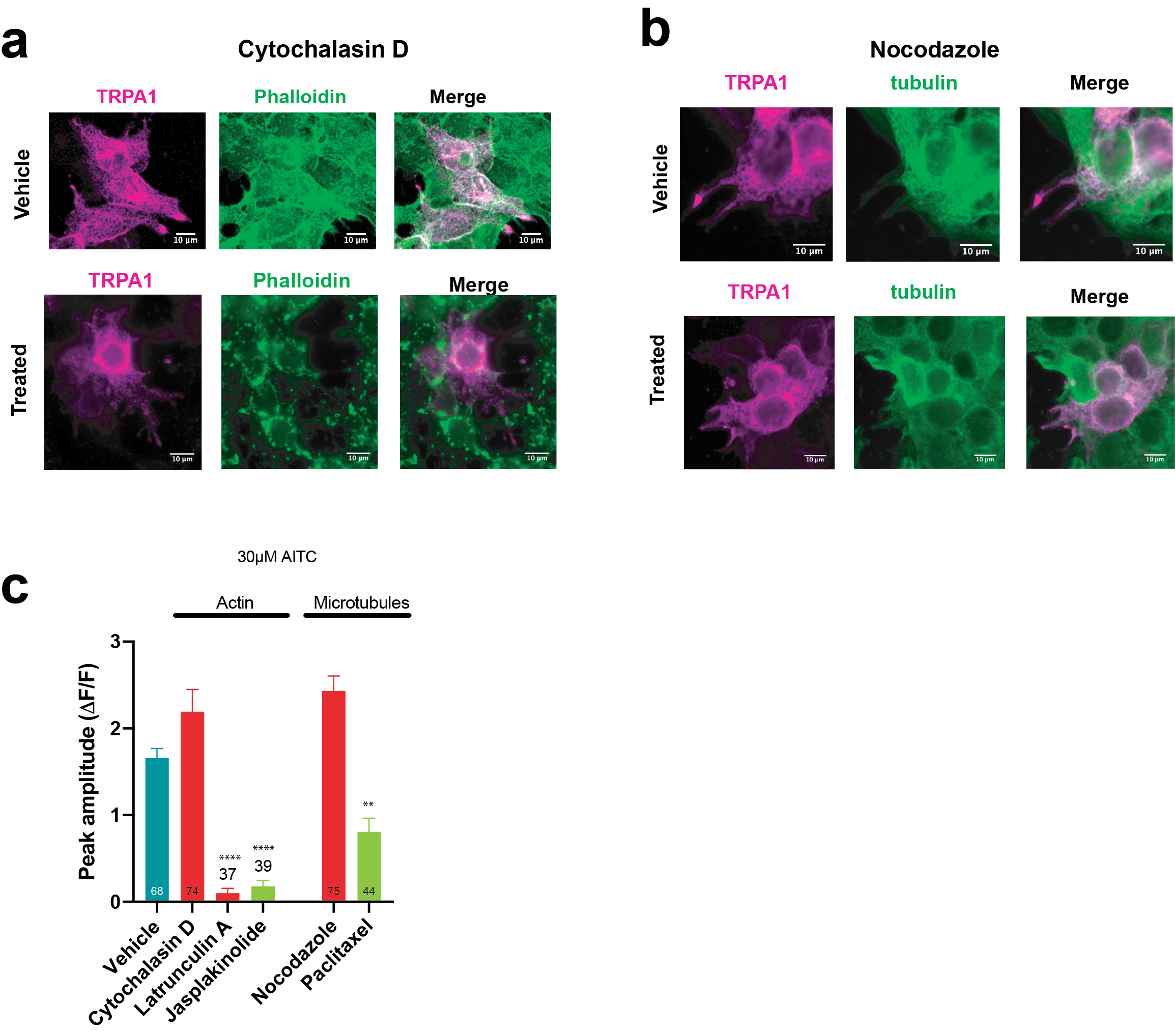

**Extended Data Fig. S5. Cytoskeletal inhibitors alter hsTRPA1 cell morphology and function.** HEK293 cells expressing hsTRPA1 have disrupted **a,** microtubules after treatment with nocodazole, **b**, actin filaments after cytochalasin-D treatment, but not vehicle controls. Microtubules are labeled using anti-alpha tubulin, while actin filaments are assessed by phalloidin staining. **c,** Treating HEK293 cells expressing hsTRPA1 with cytochalasin D or nocodazole has no significant effect on AITC responses compared to vehicle controls. In contrast, HEK293-hsTRPA responses to AITC were reduced after treatment latrunculin A and jasplakinolide and paclitaxel, presumably due to poor cell health. n = 3 coverslips/condition. Numbers of cells analyzed are shown in each bar. ** p < 0.01, **** p < 0.0001 Kruskal-Wallis rank test and Dunn’s test for multiple comparison

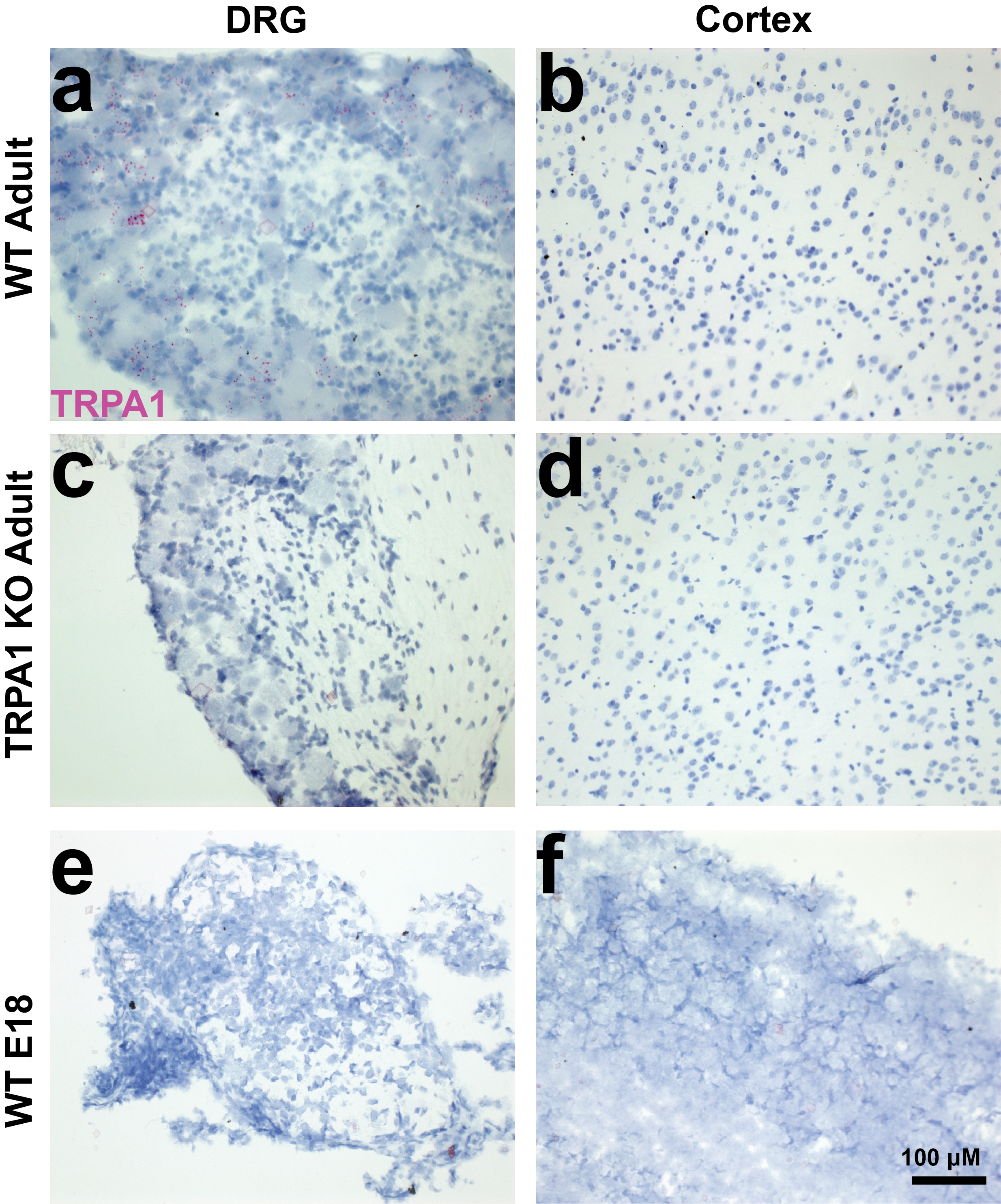

**Extended Data Fig. S6. hsTRPA1 RNA is not detected in the E18 or adult mouse cortex.** Results from a Base Scope *in situ* hybridization experiment in adult DRG and cortex taken from **(a, b)** wild-type (WT) C57Bl6/J mouse or **(c, d)** TRPA1 -/- mice as well as E18 **(e)** DRG and **(f)** cortex taken from a WT C57Bl6 embryo. Positive signal is detected as magenta puncta within cell bodies and was only detected in the adult WT DRG, as expected.

**Extended Data**
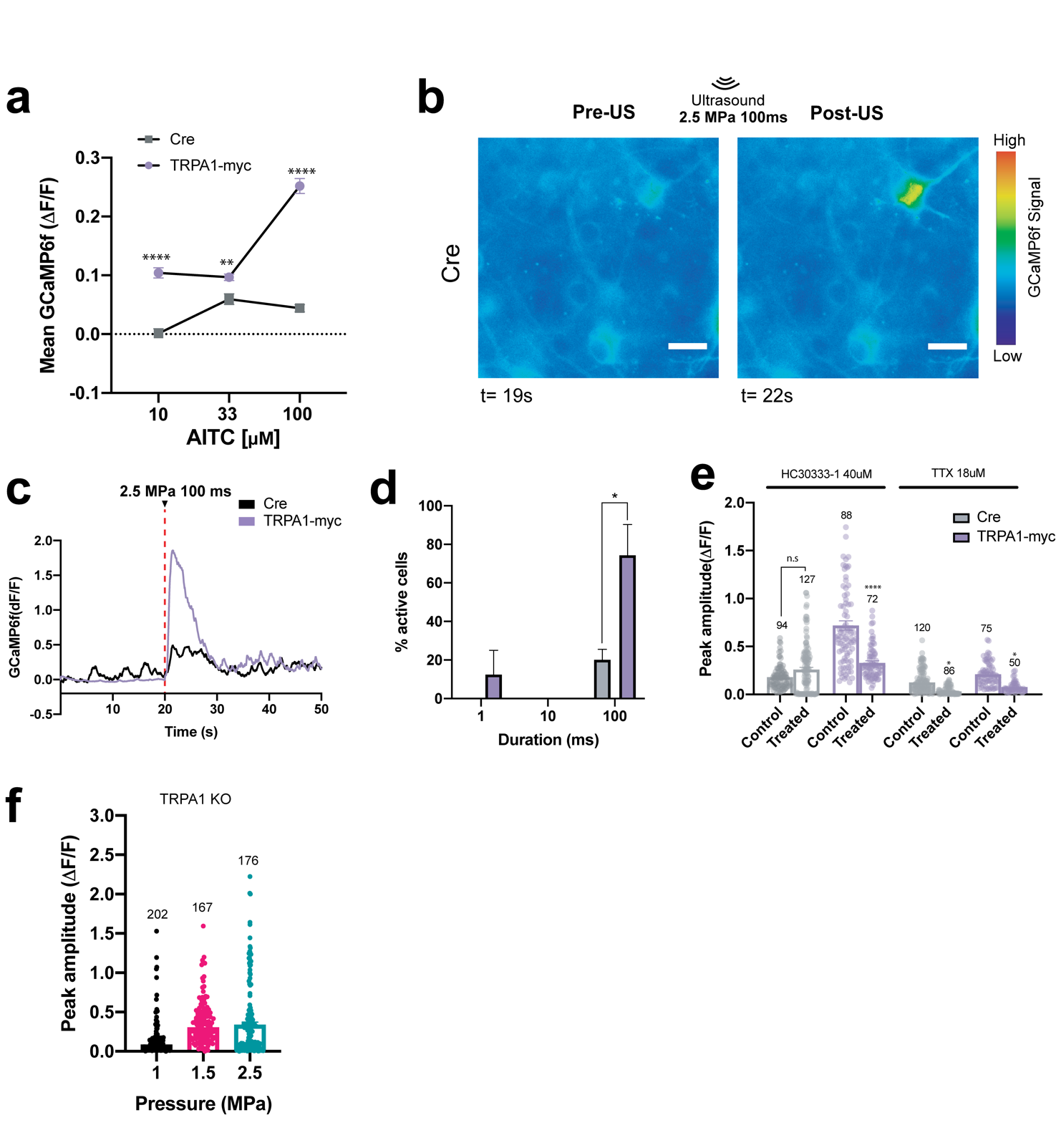
**Fig. S7. Ultrasound-evoked responses in primary neurons are independent of TRPA1**. (**a**) Dose response curve of hsTRPA1-, and Cre-control expressing neurons to AITC. n = 3 coverslips/condition. (**b**) Image showing GCaMP fluorescence in primary neurons infected with control Cre virus before and after ultrasound stimulation. (**c**) Representative traces showing magnitude of ultrasound-induced responses in representative control (Cre) or hsTRPA1-expressing neurons. (**d**) Primary neurons from TRPA1 knockout mice responded to ultrasound. n = 3 coverslips/condition. Number of cells analyzed is shown in each bar.

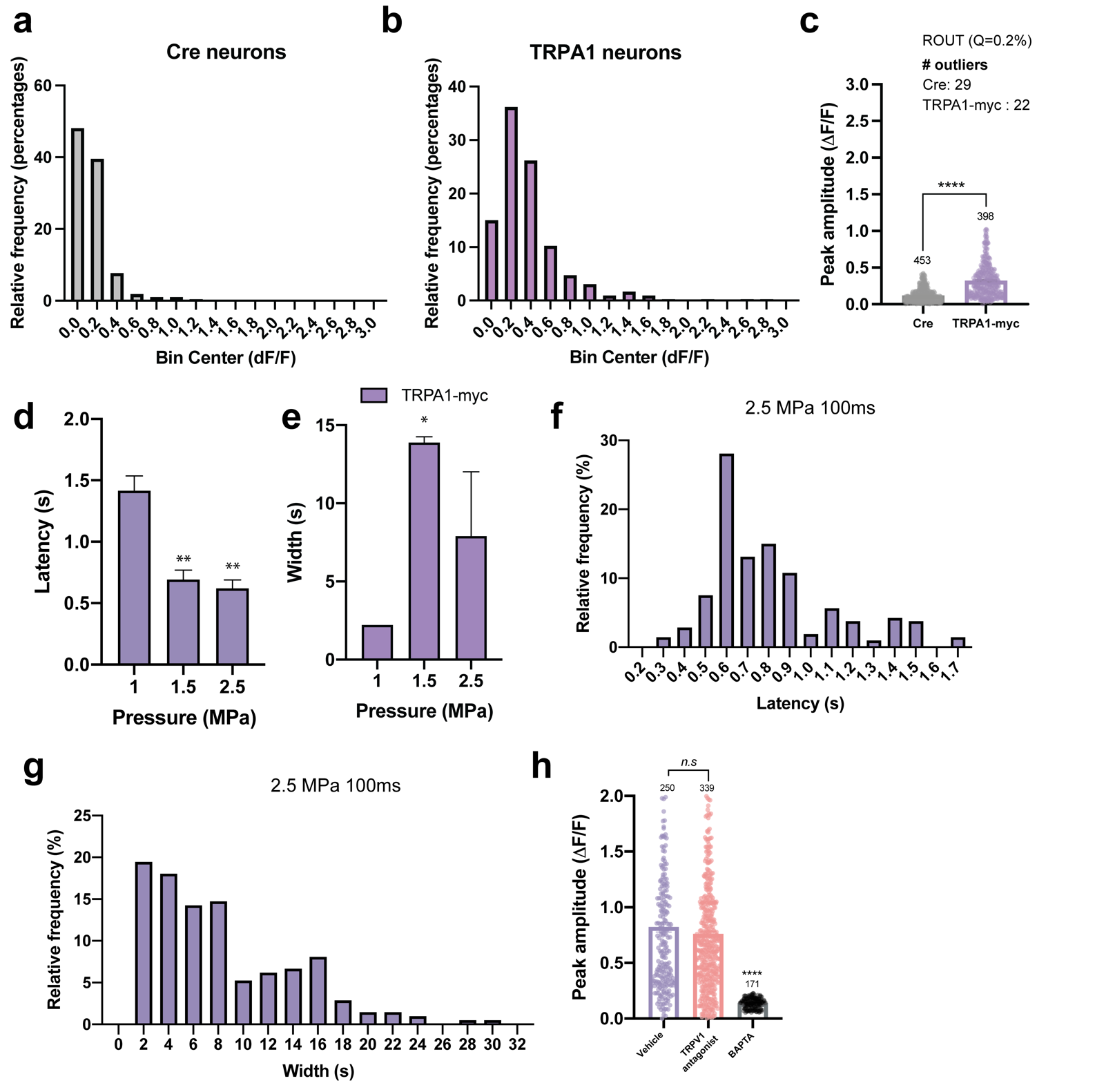

**Extended Data Fig. S8. Characterizing ultrasound responses in hsTRPA1 expressing primary neurons**. (**a**) Distribution of ultrasound responses to 100ms 2.5MPa in control and **b**, TRPA1-myc primary neurons. **c**, removing outliers reduces the maximum value observed for TRPA1-myc infected neurons but we still observe a statistically significant difference between controls and TRPA1 (pv<0.001) confirming the robustness of the effect. Plot showing **d**, time to 60% of peak response (latency) and **e**, time between 63% rise and 63% decay (response width) after ultrasound stimulation at 100 msecs and different peak negative pressures in hsTRPA1 expressing primary neurons. Plots showing distribution of **f,** latency) and **g,** response width after ultrasound stimulation in hsTRPA1 expressing neurons. (**h**) Plot showing GCaMP6f peak amplitude in hsTRPA1 expressing neurons after ultrasound stimulation and treatment with either TRPV1 antagonist (A784168, 2 μM), Calcium chelator (BAPTA, 30 μM) or vehicle (DMSO). n = 3 coverslips/condition. Numbers of cells analyzed is shown in each bar. * p<0.05, ** p<0.01 by one-way ANOVA, (**h**) ****p<0.0001, n.s, not significant p>0.05 by Kruskal-Wallis rank test and Dunn’s test for multiple comparisons.

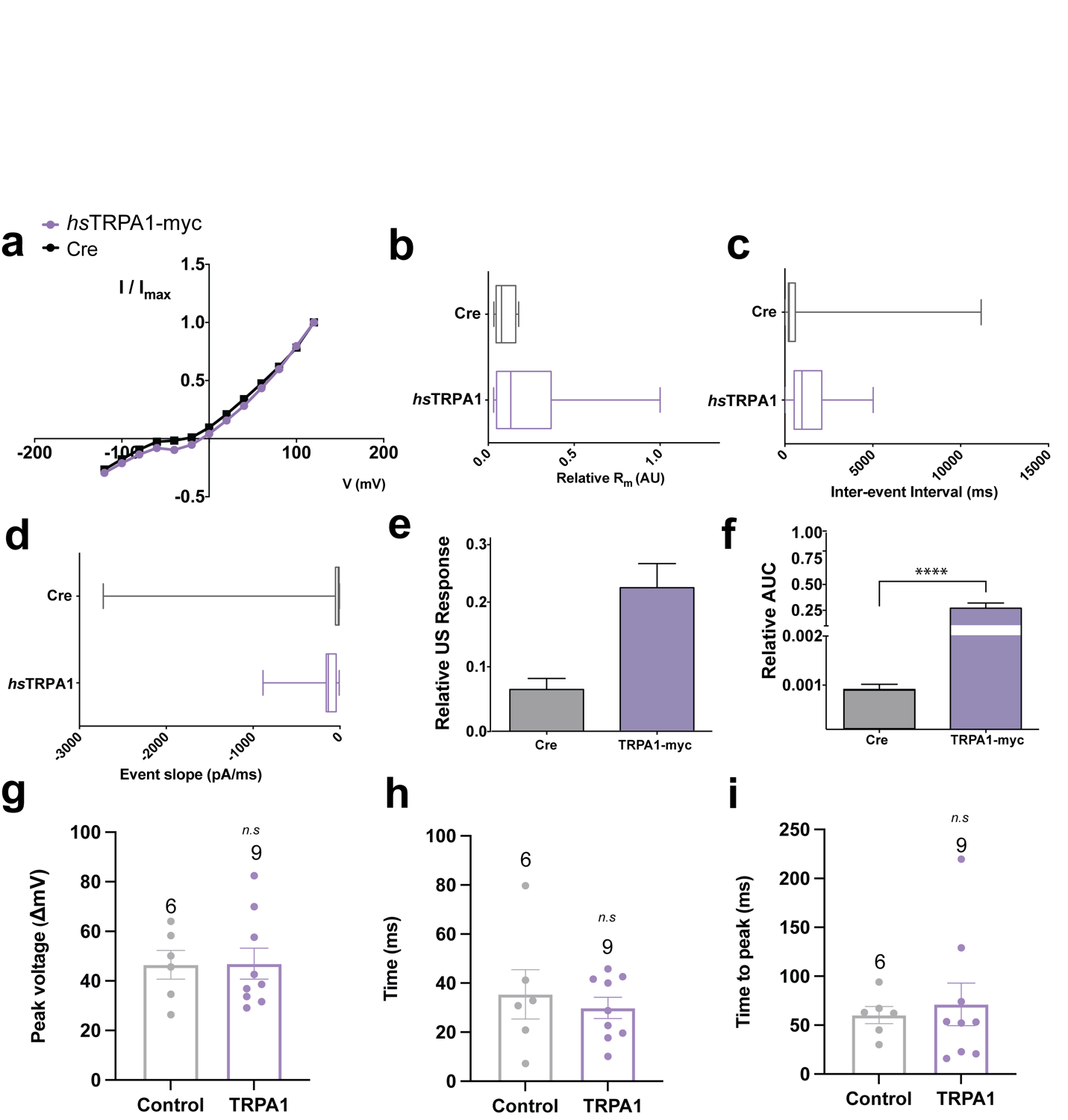

**Extended Data Fig. S9. Electrophysiological properties of primary neurons.** Functional and membrane properties are similar between TRPA1 and Cre-control infected neurons. Current-Voltage (IV) plots **a,** for AAV9-hsTRPA1 vs AAV9-Cre control primary neurons elicit similar responses. Membrane resistance can be used as a proxy for patch and recording quality **b**, similar Rm was observed for both groups. Other response characteristics including inter-event interval **c,** and response slopes **d,** were not significantly altered between TRPA1 and Cre-control infected neurons. **e,** Relative response to ultrasound was significantly increased in TRPA1-expressing neurons, as was **f,** AUC of the response. N=5 cells/group. Ultrasound induced action potentials show similar metrics both in control and TRPA1 expressing neurons, including the peak voltage (**g**), latency relative to ultrasound stimulus (**h**) and time to peak (**i**).

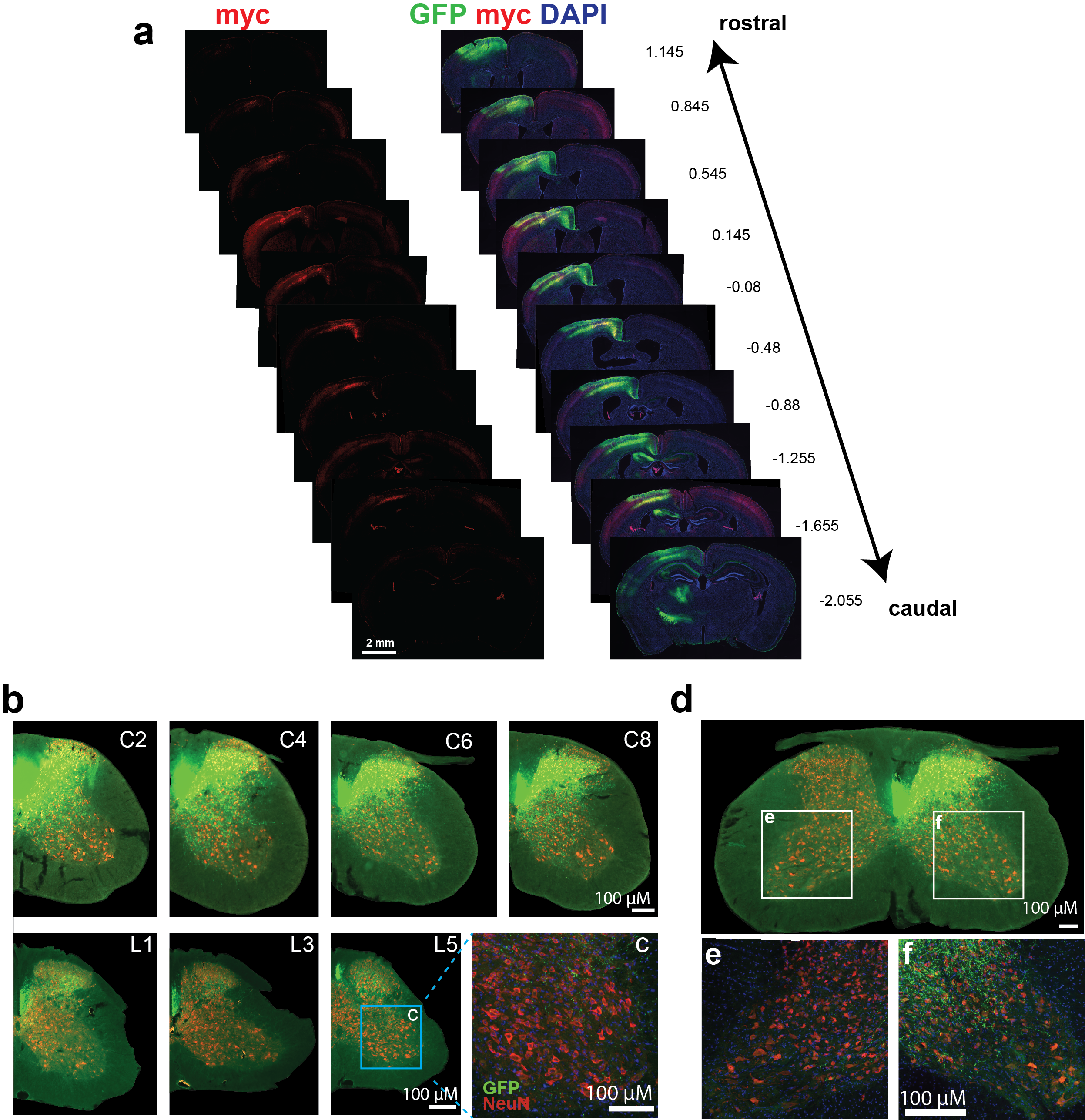

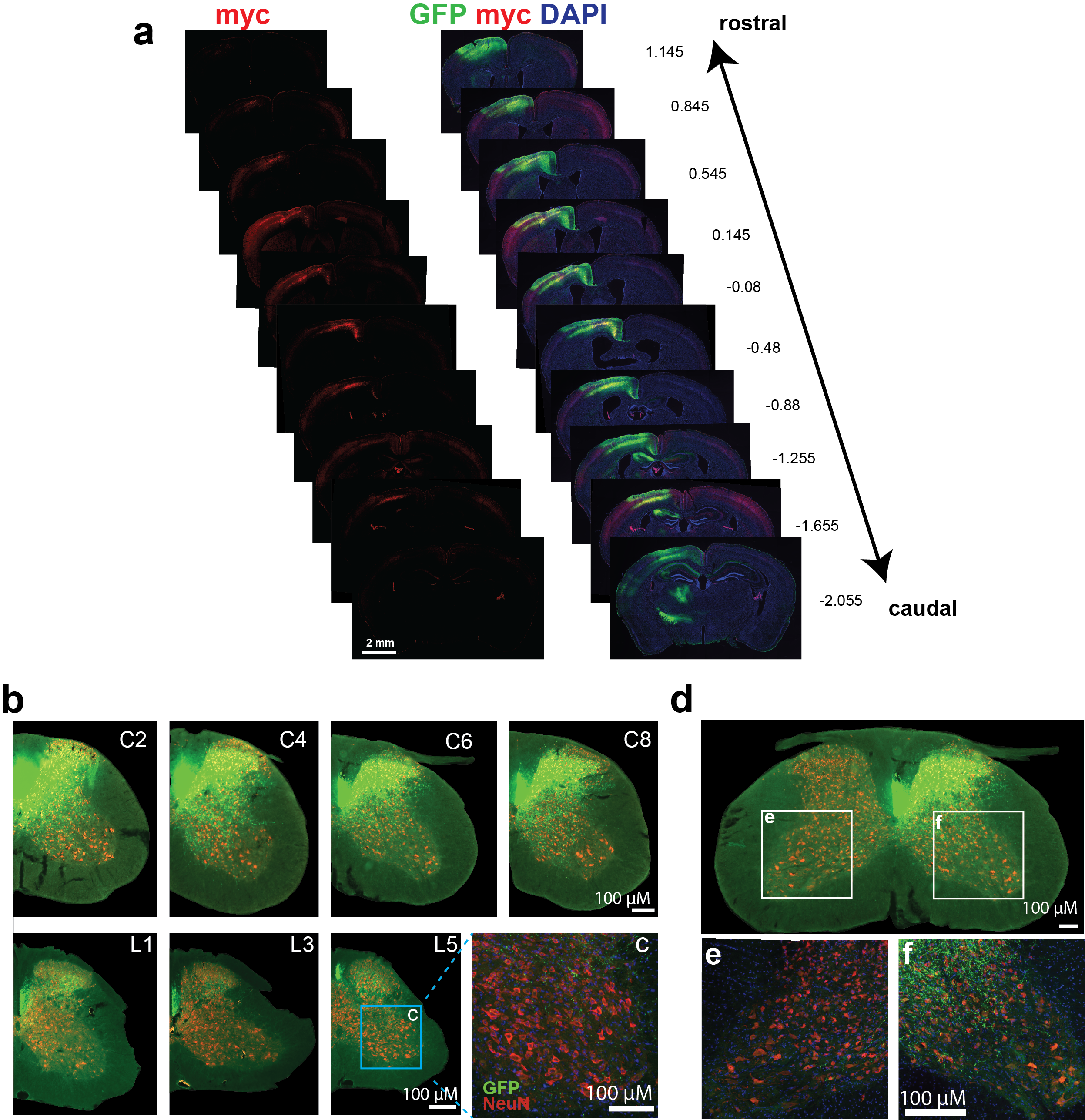

**Extended Data Fig. S10. myc-TRPA1 expresses in forelimb and hindlimb motor cortex, innervating lumbar and cervical spinal cord.** (**a**) Brain sections taken every ~350 μM were immunolabeled for myc, GFP and DAPI to evaluate the rostro caudal extent of viral expression. Approximate AP coordinates are taken from (Allen Brain Atlas(*31*)). **b**, Spinal cord sections taken every ~875 μM were immunolabeled for GFP and NeuN to evaluate the projection pattern of Npr3-Cre neurons that took up injected virus. Images are from a mouse that received co-injection of 4E13 myc-*hs*TRPA1 and 1 E12 GFP. Images were collected at 10x. **c,** A 20x confocal image of the inset from L5 showing GFP+ axons innervating the ventral horn. **d,** C6 spinal cord from the same mouse showing GFP+ axons in the **e**, ipsilateral and **f,** contralateral ventral horns.

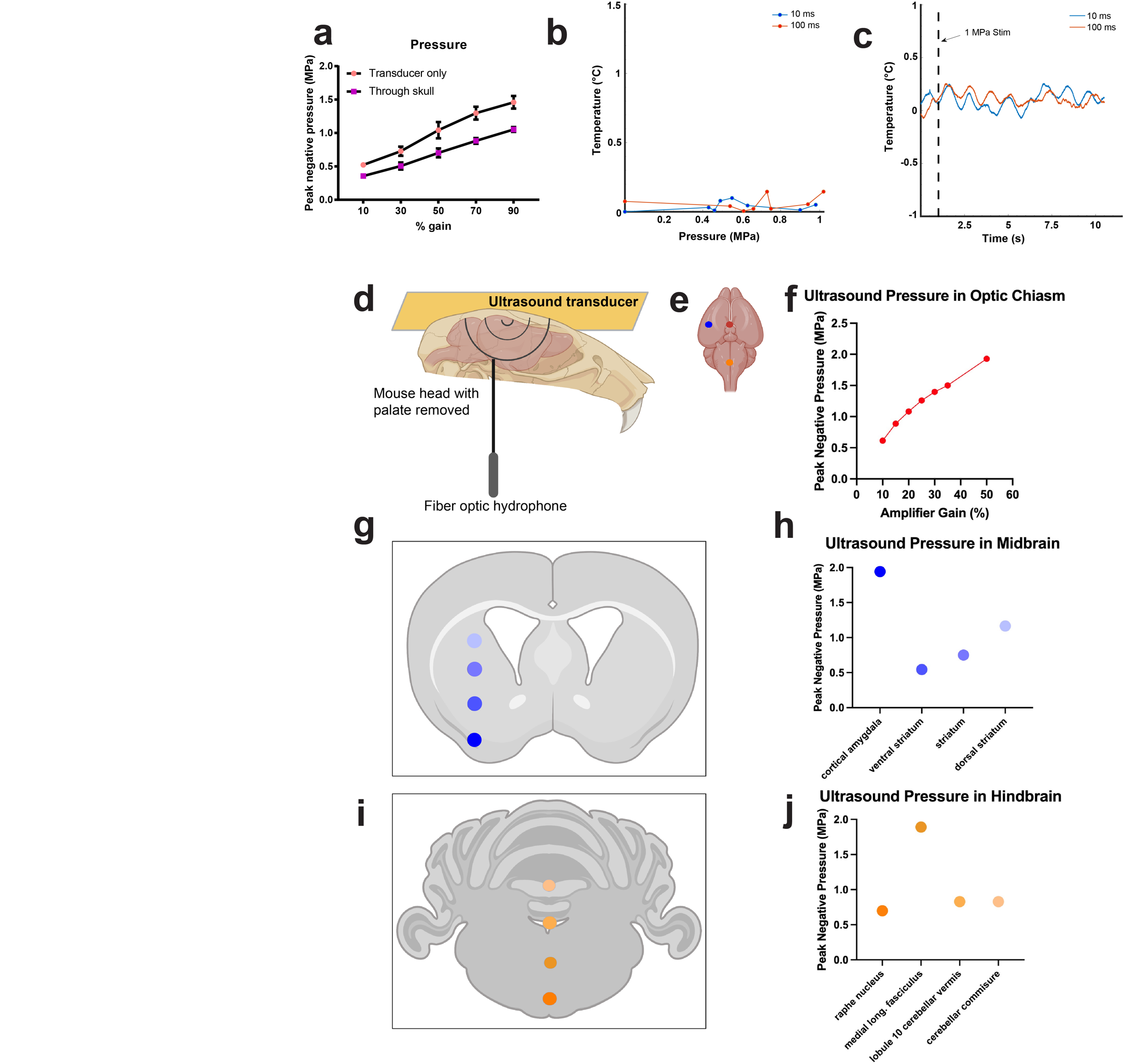

**Extended Data Fig. S11. Pressure-temperature profile of ultrasound delivery *in vivo*. a,** Pressure profile of the ultrasound transducer used for *in vivo* experiments. Peak negative pressure was measured at a consistent location relative to the face of the transducer either through ultrasound gel, or in the cortex while the ultrasound transducer was coupled to the skull with ultrasound gel. Transducer pressure output increased as a function of changing the % gain on the amplifier. **b,** Peak temperature change measured 1mm from the face of the transducer or in the cortex in response to 10 and 100ms ultrasound stimulation at increasing pressures (reported pressures are those measured within the cortex). **c**, Representative temperature traces recorded within the cortex in response to stimulation at 0.70 MPa peak negative pressure at 10 or 100 ms stimulus durations. **d**, Schematic of hydrophone recordings in ex vivo mouse brain, with skull intact and palate removed. **e**, Red dot indicates hydrophone location at optic chiasm, ventral-most part of the brain. Orange and blue dots indicate subsequent measurements at constant power and variable depth. **f**, Transducer can deliver >1.5MPa to deepest portions of the brain for sonogenetic applications. **g**, Representative midbrain coronal section, with blue dots representing hydrophone measurement locations. **h**, Ultrasound pressure delivered to midbrain, increased power can compensate for mid-range pressures. **i**, Representative hindbrain coronal section, with orange dots representing hydrophone measurement locations. **j**, Ultrasound pressure delivered to midbrain, increased power can compensate for mid-range pressures.

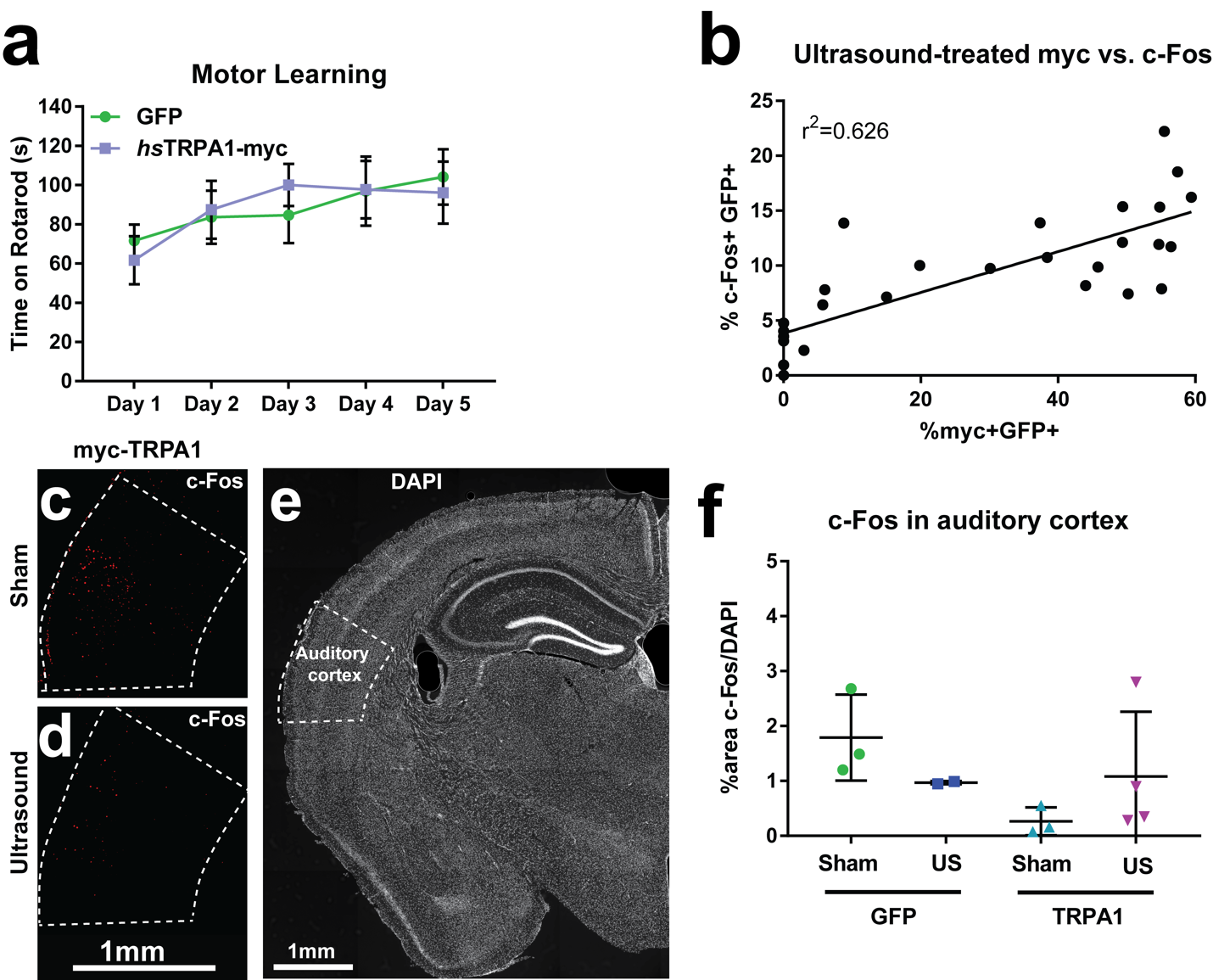

**Extended Data Fig. S12. Supplemental data from the *in vivo* experiments. a,** Rotarod performance in mice injected with 1E12 AAV9-hsyn-DIO-GFP or 1E14 AAV9-hsyn-DIO-myc:TRPA1. N=6-7 per group. No significant differences between groups by two-way ANOVA and Sidak’s multiple comparisons test. Both groups showed significant improvement in rotarod performance over the 5 days. P<0.0003 Day 5 compared to Day 1 by two-way ANOVA and Tukey’s multiple comparisons test. **b,** Correlation between %c-fos+/GFP+ neurons to %myc+/GFP+ neurons across adjacent individual sections from mice that received ultrasound treatment. R^2^=0.626. P=0<0.001. Images of c-fos in the auditory cortex from myc-TRPA1-expressing mice that received **(c)** sham stimulation or **(d)** 1hr of 100msec 1.05MPa stimulation delivered every 10 secs. **e,** Anatomical localization of auditory cortex in DAPI-labelled tissue. **f,** Quantification of % area of auditory cortex containing c-fos+ signal normalized to % area of the DAPI signal. No significant differences were detected across groups by One-way ANOVA.

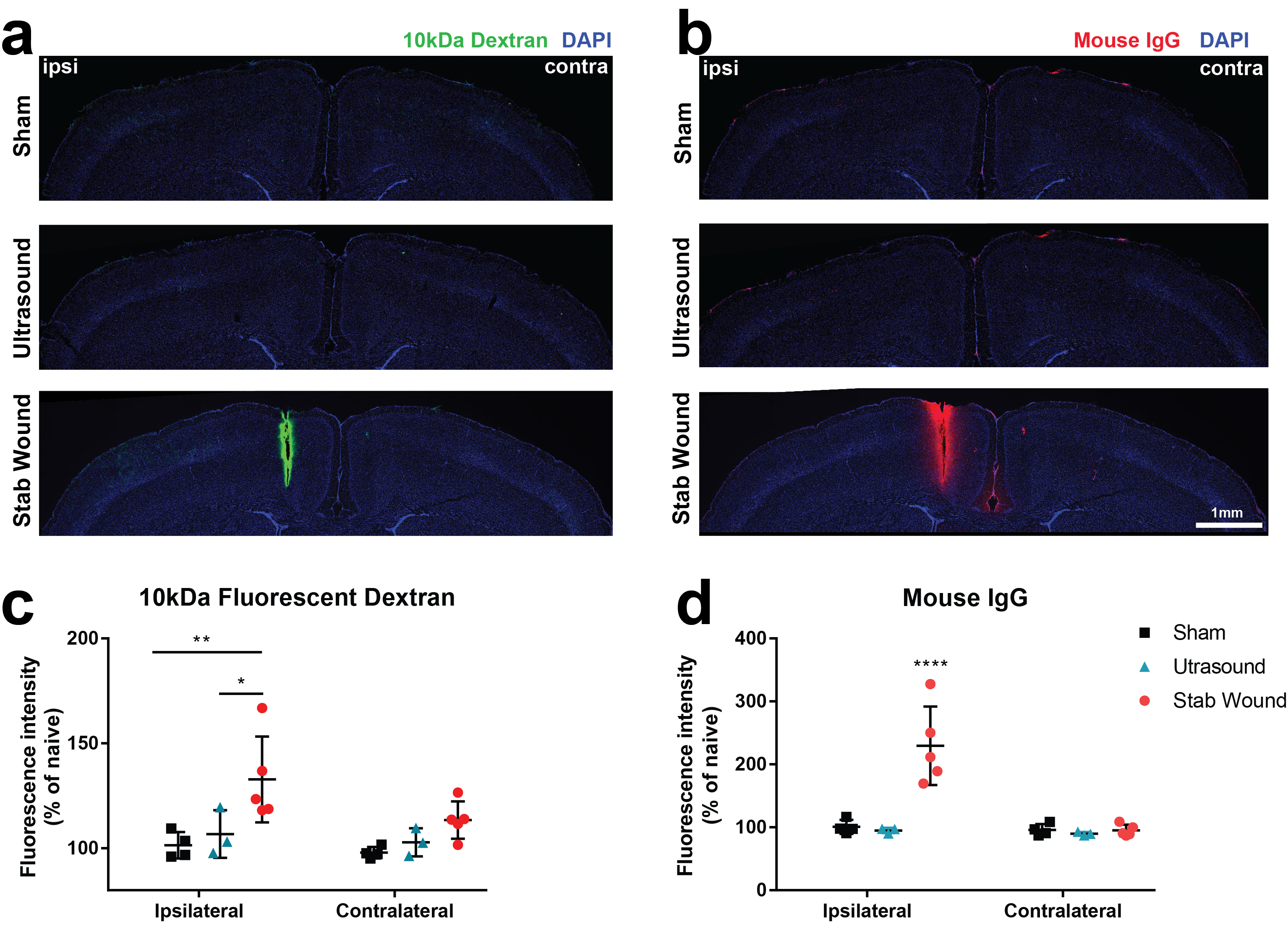

**Extended Data Fig. S13. The blood brain barrier is not disrupted by 1 hr of intermittent 100ms ultrasound delivered at 1.0 MPa.** Representative images of **a,** cortical fluorescent dextran and **b,** mouse IgG immunolabeling across conditions. **c**, Quantification of 10 kDa fluorescent dextran in each cortical hemisphere from mice that were treated with either ultrasound (100 ms, 1.0 MPa every 10 s) or sham stimulation for 1 hour or that had received a cortical stab wound condition, normalized to the cortical fluorescence of uninjected naïve mice. **d**, Quantification of mouse IgG in in each cortical hemisphere from mice that were treated with either ultrasound (100 ms, 1.0 MPa every 10 s) or sham stimulation for 1 hour or that had received a cortical stab wound condition, normalized to the cortical fluorescence of uninjected naïve mice. *p<0.05, **p<0.01, ****p<0.0001 by two-way ANOVA followed by Sidak’s multiple comparison’s test. N=3-5/group.

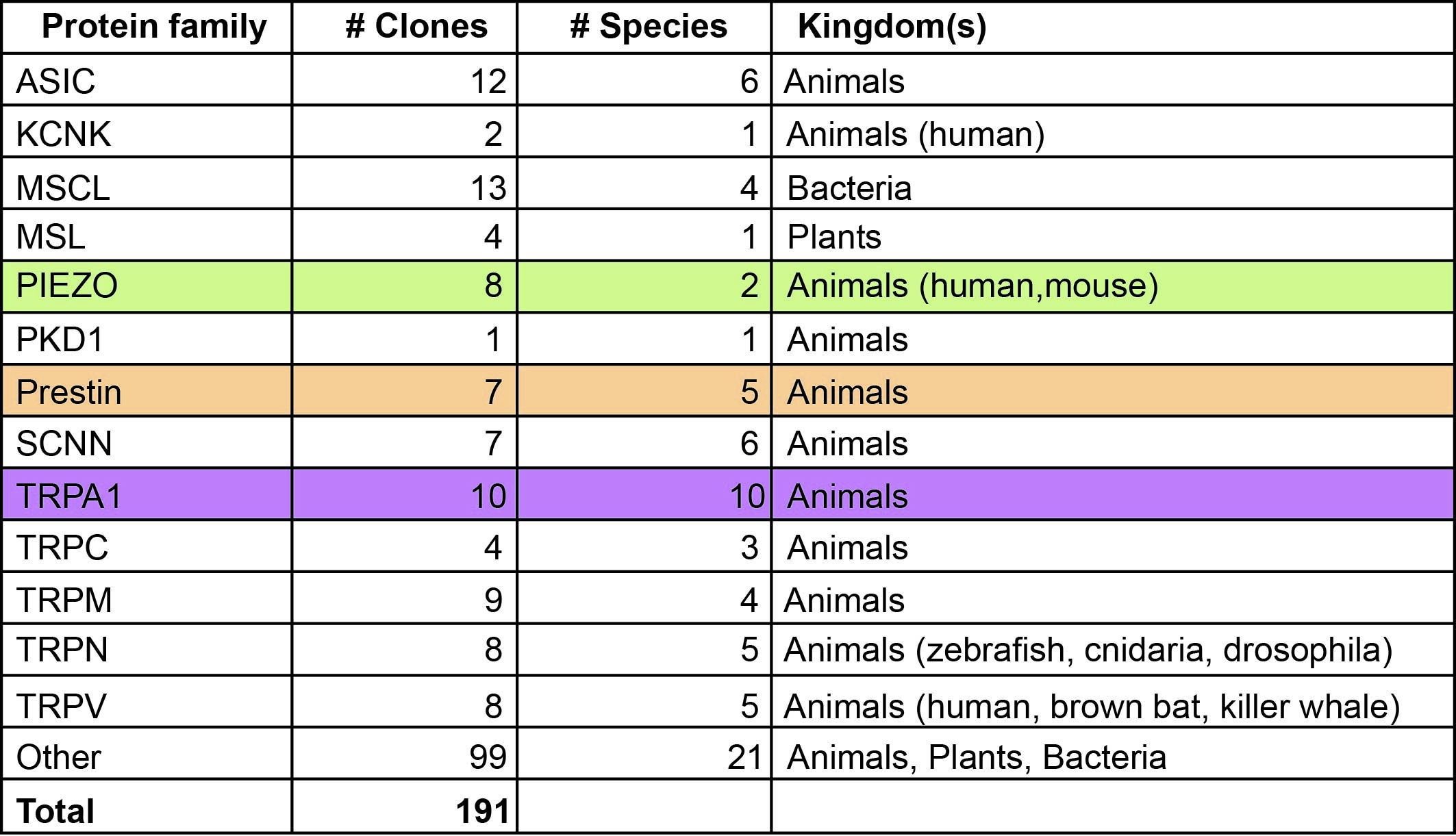

**Extended Data Table S1.** Table showing a library of 191 clones from various protein families including DEG/ENaC, K2P, TRP, ASIC, Piezo, MscS, MscL and Prestin from multiple different species.

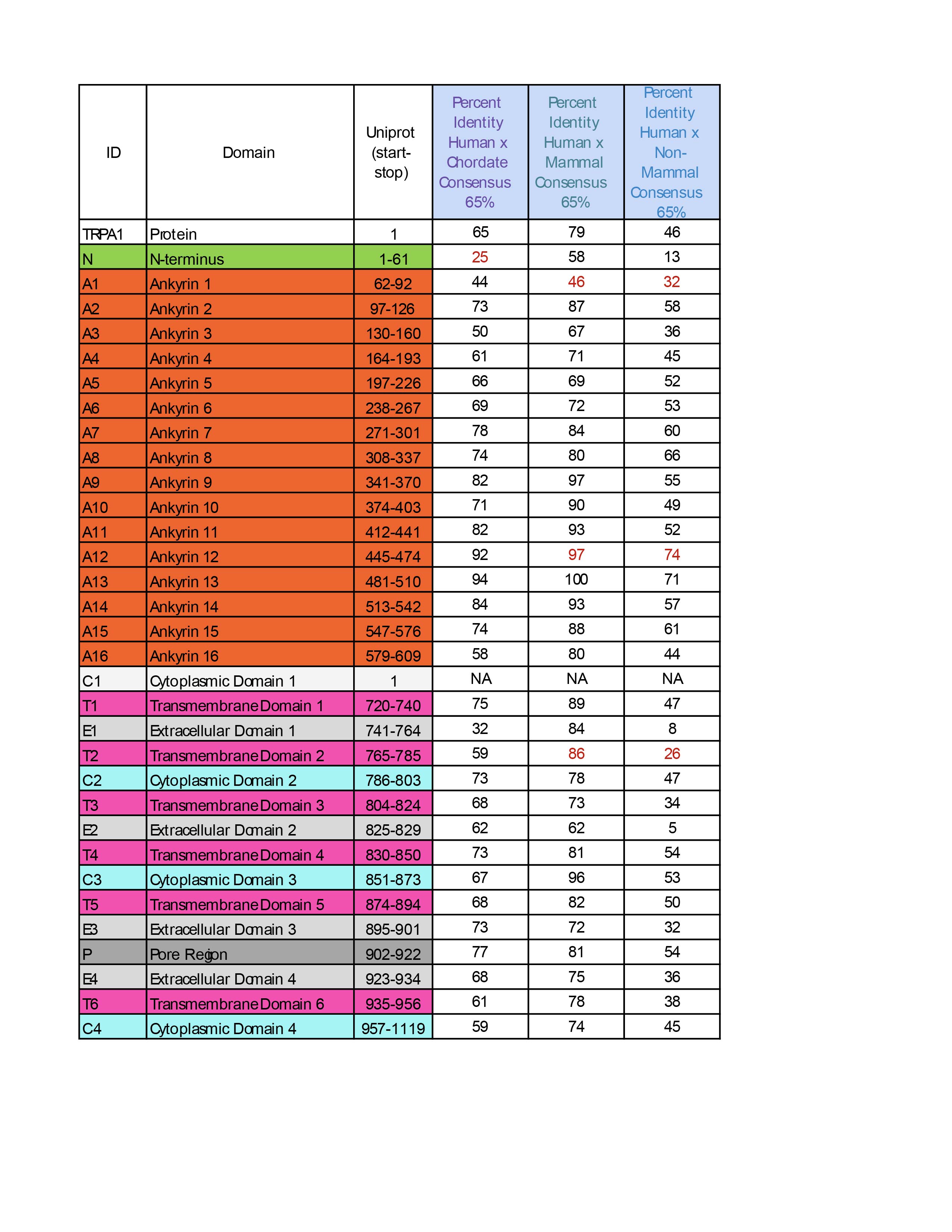

**Extended Data Table S2.** Table showing percent identity across all TRPA1 domains based on pair-wise alignment of consensus sequence for tested chordate, mammalian, and non-mammalian clades compared to human. Percent identity marked in red indicates regions that are particularly conserved or divergent between mammals and non-mammalian chordates. Threshold for consensus is bases matching to human reference in 65% of sequences in multiple sequence alignments of each clade.

**Legends for Supplementary Videos**

**Video S1. TRPA1-HEK cells respond to single ultrasound pulses at 6.91MHz**. Representative response to a single 100ms ultrasound pulse in *hs*TRPA1-expressing HEK cells. Color scale: red, high GCaMP6f ΔF/F signal, blue, low GCaMP6f ΔF/F. Scale bar 20 μm. Time scale is in seconds. Stimulation parameters: 100ms 2.5 MPa 6.91MHz delivered at t = 20 s.

**Video S2. HEK cells are normally insensitive to ultrasound at 6.91MHz**. Lack of response to a single 100ms ultrasound pulse in dTomato-expressing HEK cells, as assessed by calcium imaging. Color scale: red, high GCaMP6f ΔF/F signal, blue, low GCaMP6f ΔF/F. Scale bar 20 μm. Time scale is in seconds. Stimulation parameters: 100ms 2.5 MPa 6.91MHz delivered at t = 20 s.

**Video S3. TRPA1 increases sensitivity to 6.91MHz ultrasound in mouse primary neurons *in vitro.*** Representative response to a single 100ms ultrasound pulse in mouse cortical primary neurons (DIV10-11) that were infected with AAV9-hSyn-DIO-*hs*TRPA1 and AAV9-Cre. Color scale: red, high GCaMP6f ΔF/F signal, blue, low GCaMP6f ΔF/F. Scale bar 20 μm. Time scale is in seconds. Stimulation parameters: 100 ms 2.5 MPa 6.91 MHz delivered at t = 20 s.

**Video S4. Mouse primary neurons intrinsically show modest responses to 6.91MHz ultrasound.** Representative response to a single 100ms ultrasound pulse in control mouse cortical primary neurons (DIV10-11) that were infected with only AAV9-Cre. Color scale: red, high GCaMP6f ΔF/F signal, blue, low GCaMP6f ΔF/F. Scale bar 20 μm. Time scale is in seconds. Stimulation parameters: 100 ms 2.5 MPa 6.91 MHz delivered at t = 20s.

**Video S5. *hs*TRPA1 expression allows robust repeated ultrasound stimulation in mouse primary neurons in vitro**. This video shows a representative response to repetitive100ms ultrasound pulses delivered every 10s (0.1Hz) in mouse cortical primary neurons (DIV10-11) that were infected with AAV9-hSyn-DIO-TRPA1 and AAV9-Cre. Color scale: red, high GCaMP6f ΔF/F signal, blue, low GCaMP6f ΔF/F. Scale bar 20 μm. Time scale is in seconds. Stimulation parameters: 100 ms 2.5 MPa 6.91MHz.

**Video S6. *hs*TRPA1 expression allows for ultrasound-evoked contralateral hindlimb movement.** This video shows a representative right hindlimb (contralateral to injection site) response to repetitive 0.88 MPa ultrasound delivered for 10 ms or 100 ms in an Npr3-cre mouse expressing *hs*TRPA1 in the left motor cortex. The red LED and “Ultrasound” text indicate when ultrasound is on. No left limb movements were observed.
